# Comparative genomics and transcriptomic response to root exudates of six rice root-associated *Burkholderia sensu lato* species

**DOI:** 10.1101/2022.10.04.510755

**Authors:** Adrian Wallner, Agnieszka Klonowska, Ludivine Guigard, Eoghan King, Isabelle Rimbault, Eddy Ngonkeu, Phuong Nguyen, Gilles Béna, Lionel Moulin

**Author notes:** Corresponding author: Lionel Moulin. Present address: Unité Résistance induite et Bio-protection des plantes, University of Reims Champagne-Ardenne, Reims, France (Adrian Wallner). Present address: Centro de Biotecnología y Genómica de Plantas (CBGP), Universidad Politécnica de Madrid (UPM)–Instituto Nacional de Investigación y Tecnología Agraria y Alimentación (INIA/CSIC), Campus de Montegancedo, Pozuelo de Alarcón (Madrid), Spain.

## Abstract

Beyond being a reliable nutrient provider, some bacteria will perceive the plant as a potential host and undertake root colonization leading to mutualistic or parasitic interactions. Bacteria of the *Burkholderia* and *Paraburkholderia* genera are frequently found in the rhizosphere of rice. While the latter are often described as plant growth promoting species, *Burkholderia* are often studied for their human opportunistic traits. Here, we used root exudate stimulation on three *Burkholderia* and three *Paraburkholderia* strains isolated from rice roots to characterize their preliminary adaptation to the rice host at the transcriptomic level. Instead of the awaited genus-dependent adaptation, we observed a strongly species-specific response for all tested strains. While all bacteria originate from the rice environment, there are great disparities in their levels of adaptation following the sensing of root exudates. We further report the shared major functions that were differentially regulated in this early step of bacterial adaptation to plant colonization, including amino acids and putrescine metabolism, the Entner-Doudoroff (ED) pathway as well as cyclic diguanylate monophosphate (c-di-GMP) cycling.

## Background

Diverse species of *Burkholderia* and *Paraburkholderia* have been repeatedly isolated from healthy rice roots (Govindarajan et al., 2008; Mattos et al., 2008; Wang et al., 2016a; Kang et al., 2017; Arriel-Elias et al., 2019; Song et al., 2019; Hameed et al., 2020). Their central role in rice ecology is further suggested by several population analyses placing *Burkholderia* and *Paraburkholderia* as being among the predominant taxa found in the rice rhizosphere with some specimens acting as microbiome structuring hubs (Ikeda et al., 2014; Hardoim, 2015; de Souza et al., 2015; Yu et al., 2018).

Both *Burkholderia* and *Paraburkholderia* are described as efficient rhizosphere colonizers and can achieve plant growth promotion through diverse strategies including nutrient solubilization, phytohormone production and interference, siderophore production as well as enhancing the plants tolerance to abiotic stresses. Several *Paraburkholderia* species are free living nitrogen fixers and some induce nitrogen-fixing nodules on legume plants (Dall’Agnol et al., 2017; Estrada-de los Santos et al., 2018). Further *Paraburkholderia* species are root endophytes and enhance the growth of a number of major crops such as corn, rice and sugarcane (Perin et al., 2006; Govindarajan et al., 2008; Mattos et al., 2008; Mitter et al., 2013). While several *Burkholderia* species e.g., *B. glumae, B. gladioli, B. plantari*, are responsible for serious diseases in rice crops (Kim et al., 2021), others seem to have an intrinsic ability to colonize roots which is independent of plant genotype and do not necessarily cause disease (Vidal-Quist et al., 2014). *B. cenocepacia* or *B. vietnamiensis* for instance have been observed to cause plant tissue water-soaking in onion but have otherwise been isolated from healthy plants (Engledow et al., 2004; Jacobs et al., 2008). While *Burkholderia* strains are mostly reported for their biocontrol efficiency, some achieve plant growth promotion through direct hormonal stimulation and nutrition (Trân Van et al., 2000; Araújo et al., 2016; Batista et al., 2016; Wang et al., 2016b; Mullins et al., 2019; Chávez- Ramírez et al., 2020). Genome mining approaches underlined the importance of secondary metabolites in *Burkholderia* species, including an important diversity of antimicrobial metabolites (Navarro et al., 2019). However, in contrast to *Paraburkholderia*, several *Burkholderia* species were found to colonize animal hosts in addition to plants, causing opportunistic infections and severe pathologies in humans (Galyov et al., 2010; Sousa et al., 2011).

Here, we investigated the early strategies directing plant colonization at the onset of the interaction between rhizobacteria and their host which is mediated by the perception of root exudates (RE). Through root exudation, plants can use up to 20-40% of their photosynthetically fixed carbon to actively shape their microbiome using chemical attractants, selective carbon sources and diverse restrictive metabolites (Badri and Vivanco, 2009). Primarily genes involved in chemotaxis and motility, but also nutrient transport and metabolism are required for an efficient response to exudates. Still, as exemplified by several past studies, a bacterium’s response to exudates is highly host dependent. For instance, *Pseudomonas aeruginosa* was shown to have a varying transcriptomic reaction to RE originating from two different sugar beet cultivars (Mark et al., 2005). The perception of RE can induce major metabolic adaptation in bacteria. Rice RE were found to be responsible for the differential expression of 4,4% genes in the genome of *Azoarcus* sp. BH72 (Shidore et al., 2012). More significant, a legume nodulating strain of *Paraburkholderia phymatum* differentially regulated 19.2% of its genome in response to RE of *Mimosa pudica* (Klonowska et al., 2018). Upon inoculation with maize RE, *Bacillus amyloliquefaciens* displays a two- stage response. At 24h post-treatment, the bacteria directs its metabolism towards cellular proliferation while at 48h post-treatment, most differentially expressed (DE) genes are down- regulated and production of extracellular matrix is favored (Fan et al., 2012; Zhang et al., 2015). Typically, the bacterial reaction to RE involves metabolic genes for sugar and amino- acid import and catabolism. However, functions involved at later steps of the interaction such as type 3 secretion are modulated as well upon RE perception (Hassan and Mathesius, 2012; Klonowska et al., 2018).

The present work focuses on six rice root-isolated strains evenly distributed between *Burkholderia* and *Paraburkholderia* genera. Beside the two model bacteria for plant growth promotion *Burkholderia vietnamiensis* LMG 10929 (BvLMG10929) and *Paraburkholderia kururiensis* M130 (PkM130), we analyzed four new rice-isolated bacterial strains, for which we provide whole-genome sequences, as well as an assessment of their rice root colonization capacity. We hypothesized that the genetic frontier between the two bacterial taxa would influence and differentiate their adaptation strategies towards plant colonization in a genus specific manner. With this in mind, we aimed at revealing each strain’s adaptation signatures to the plant environment by exploring their gene expression profiles when sensing rice root exudates using a transcriptomics approach. Considering the results presented here, no distinction can be made between bacteria according to their genus as each strain displayed a specific reaction upon plant root exudate sensing, with no significant pattern clustering con- generic bacteria, while the regulation of several metabolic functions was shared among all bacteria.

## Results

### Genomic analyses of new *Burkholderia* and *Paraburkholderia* species

The newly isolated strains described in this study (Table 1) were compared to genomic databases of *Burkholderia sensu lato* (*s.l.*) and the genomic distance to their closest relative was computed (Figure 1). Whole genome comparisons using average nucleotide identity (ANI) estimations showed a 98.52% score (80.14% sequence coverage) when comparing strain ABIP441 (hereafter BrABIP441) to the proposed species *Burkholderia reimsis* sp. nov. BE51^T^ (Esmaeel et al., 2018), lying above the species threshold (> 95% identity on more than 70% cover). We had previously proposed that ABIP444 (hereafter BoABIP444) belongs to a novel species, *B. servocepacia* sp. nov., closely related to *Burkholderia cenocepacia* (Wallner et al., 2019). A recent publication updated our proposal to validate this separate species as *B. orbicola,* with TatI-371 as the type strain (Morales-Ruíz et al., 2022). Strain ABIP659 (hereafter PspABIP659) displays no determining similarity to any described *Burkholderia s.l.* species. ANI analyses suggest that it is most closely related to *Paraburkholderia tropica* without exceeding the required threshold to infer conspecificity (91.86% identity and 76.0% sequence coverage to *P. tropica* LMG 22274^T^). At the genome level, ABIP630 (hereafter PsABIP630) shares 98.5% ANI (>70% aligned) with *P. sabiae* LMG24235^T^ and can thus be considered as belonging to this species.

**Figure 1.**
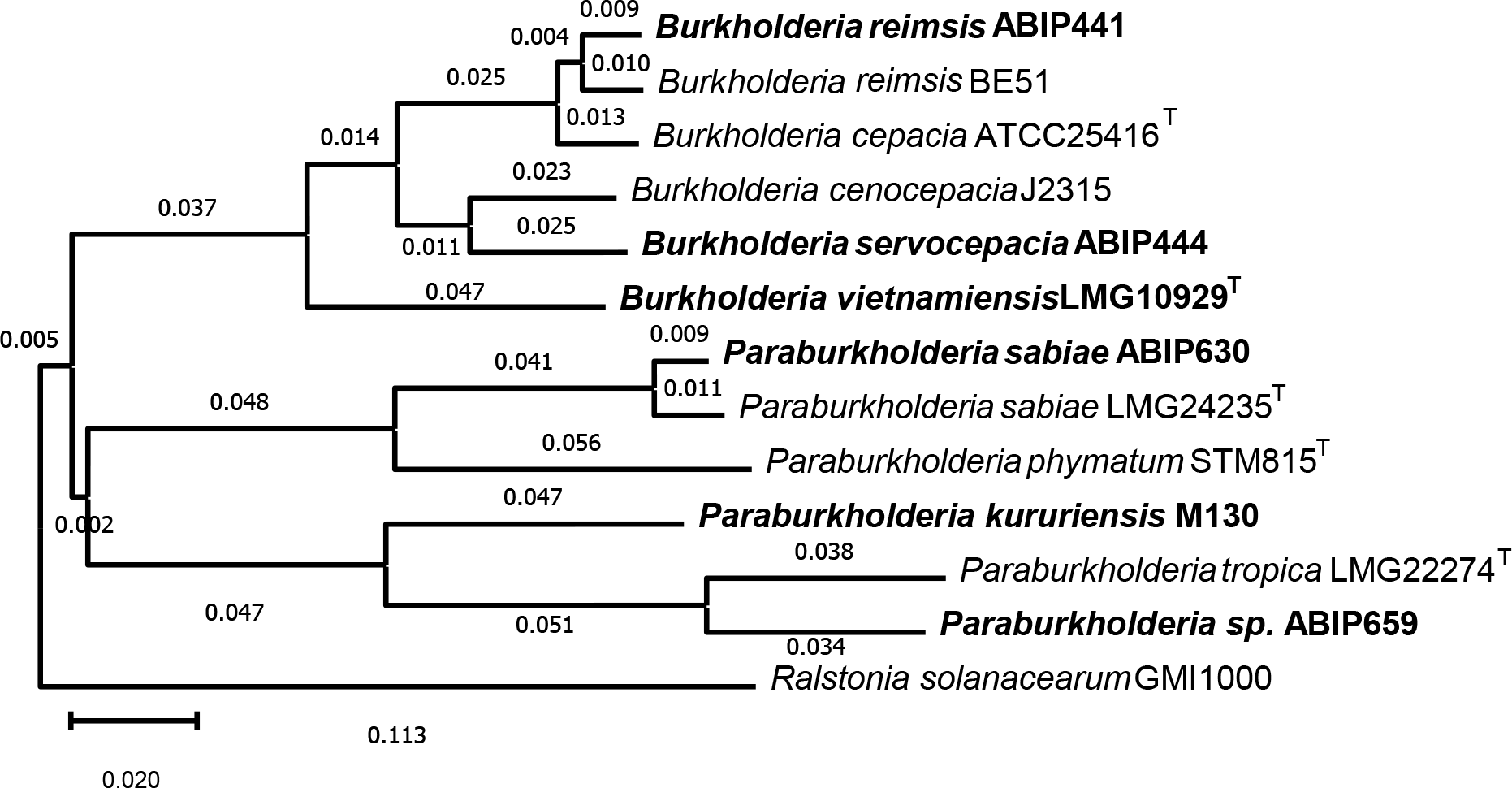
Genome clustering of studied strains (in bold) with their closest species neighbors. The tree was constructed using the Genome clustering tool of the Microscope plateform (https://mage.genoscope.cns.fr/microscope/). Distances between whole genomes was computed using Mash (Ondov et al., 2016) and plotted using the rapid Neighbor-Joining method (Simonsen et al., 2008). The displayed distances are correlated to average nucleotide identity (ANI) such as D ≈ 1-ANI. Genbank genome accession numbers of strains under study are indicated in Table 1.

**Table 1.** List of the six strains used in this study & their genomic characteristics.

|  | *B. reimsis*<br>ABIP441 | *B. orbicola*<br>ABIP444 | *B. vietnamiensis*<br>LMG10929 | *P. sabiae*<br>ABIP630 | *P. sp.*<br>ABIP659 | *P. kururiensis*<br>M130 |
| --- | --- | --- | --- | --- | --- | --- |
| Genbank | GCA_914492715 | GCA_905184115 | GCA_000959445 | GCA_914484815 | GCA_914484835 | GCA_000341045 |
| Genome size (bp) | 7845349 | 7409039 | 6930496 | 8354439 | 8573543 | 7129657 |
| Contigs | 311 | 292 | 3 Chromosomes,<br>1 Plasmid | 84 | 122 | 9 |
| Genomic objects | 8330 | 7822 | 6908 | 8239 | 8395 | 6524 |
| CDSs | 7881 | 7402 | 6416 | 7779 | 7934 | 6116 |
| RNA genes | 101 | 96 | 87 | 93 | 99 | 88 |
| Phage-associated<br>genes | 61 | 79 | 10 | 14 | 52 | 33 |
| GC% | 66.59 | 66.72 | 65.07 | 62.48 | 64.95 | 65.02 |
| **Isolation** |  |  |  |  |  |  |
| Country | Cameroon | Cameroon | Vietnam | Cameroon | Vietnam | Brazil |
| Compartment | Rhizosphere | Rhizosphere | Rhizosphere | Rhizosphere | Rhizosphere | Endosphere |
| Reference | This study | (Wallner et al.,<br>2019) | (Gillis et al.,<br>1995) | This study | This study | (Baldani et al.,<br>1997) |

### All isolates demonstrate robust rice colonization capacity

As all strains were isolated from the rice rhizosphere (Table 1), we validated their ability to colonize this environment *de novo* (Figure 2). Plants growing in a sterile hydroponic environment were inoculated with each strain individually and the root-associated strains were enumerated by plate counting after 3, 7 and 14 days. Compared to the well investigated BvLMG10929 and PkM130, all strains show successful root attachment and colonization. Most bacteria reach their highest density at 7 days post inoculation (dpi) and the ratio of bacterial cells to root mass then decreases. The three *Burkholderia spp.* all have very similar colonization efficiencies and development profiles over time. *Paraburkholderia spp.* in turn, have more individual colonization approaches. PsABIP630 displays the strongest abundance on roots while PspABIP659 is the overall weakest colonizer but has the greatest dynamic in colonizing populations. Indeed, PspABIP659 shows a striking 6-fold burst in population from 3 dpi to 7 dpi. For all strains, we evaluated the impact of inoculation on *Oryza sativa* cv. Nipponbare growth (see Methods). No impact was observed on root and leaf weight at 60 dpi compared to the mock treatment (Supplementary Figure S1). This is consistent with results from similar treatments using PkM130 and BvLMG10929 (King et al., 2019).

**Figure 2.**
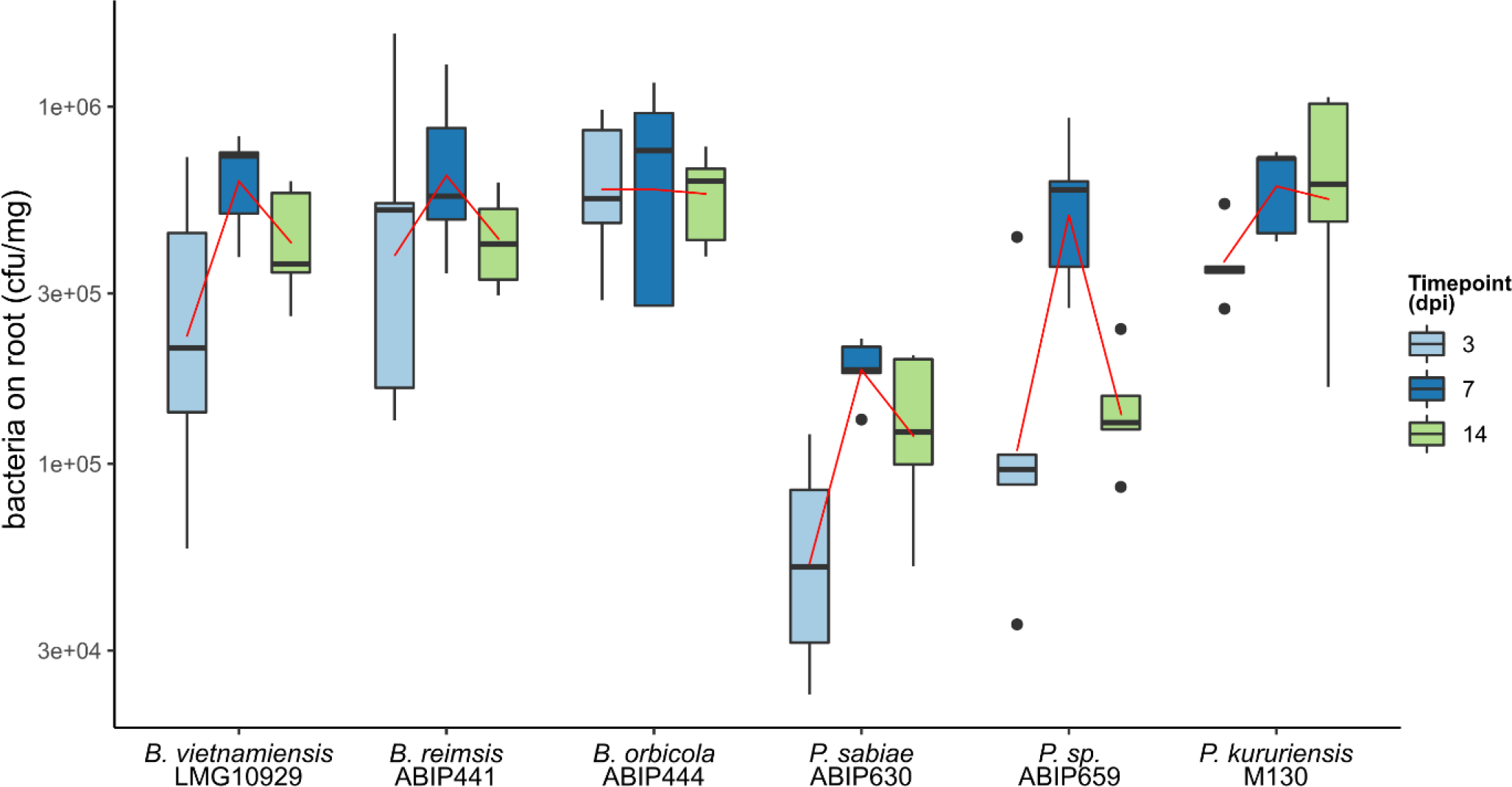
Plant colonization efficiencies for six *Burkholderia* and *Paraburkholderia* strains. Nipponbare rice seedlings were inoculated at 5 days post-germination with 1.10^8^ bacterial cells and grown in a hydroponic medium. Total roots were sampled at different time points (5 replicates each) and the attached bacterial population was enumerated. The trendlines (red) were inferred using the non-parametric regression LOESS method (Cleveland et al., 2017).

### Paraburkholderia and Burkholderia species share 2443 genes

An orthology search was performed to infer levels of similarity between the three *Paraburkholderia* strains and the three *Burkholderia* strains (Figure 3). While all genomes are of relatively large size compared to their Proteobacteria congeners (Table 1), there are some discrepancies between them. The two rice-growth promoting model endophytes PkM130 and BvLMG10929 have genomes of comparable size which are however the smallest among the cohort analyzed here. The four remaining species have all been isolated from the rice rhizosphere, and their genomes are considerably larger (15-20%) than those of the endophytic PkM130 and BvLMG10929.

**Figure 3.**
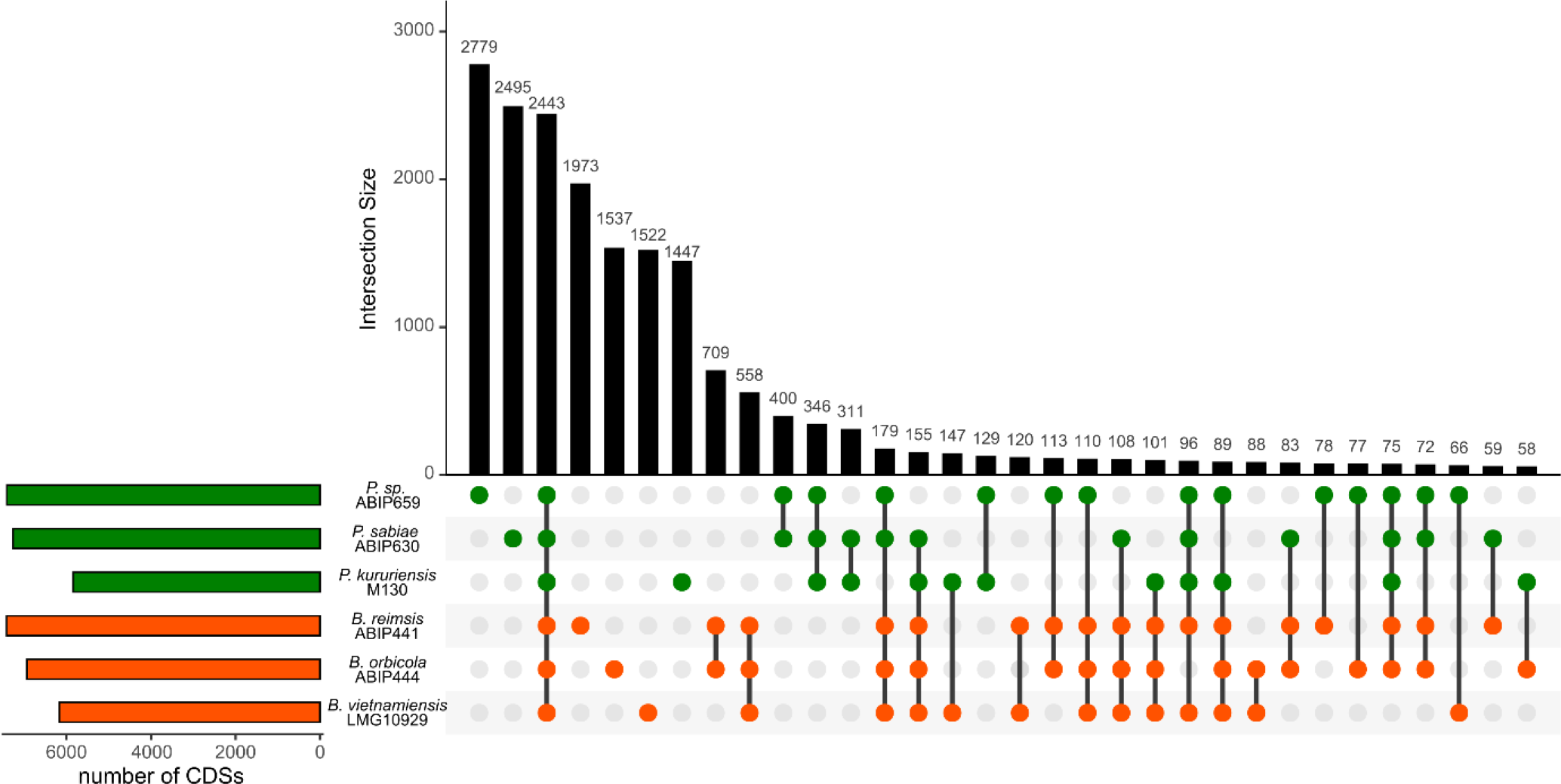
UpSet plot of gene conservation between six Burkholderia s.l. strains. Single dots are representative of the specific genome of each strain. Linked dots represent the conserved genes among the connected strains. All combinations totaling less than 50 conserved genes are not shown. Gene duplicates were excluded from this analysis.

Altogether, 2443 genes are conserved among the six strains, representing 34% to 44% of their total gene sets (Figure 3). Still, given the present strain selection, PspABIP659 and PsABIP630 are best defined by their specific genome which is larger than any other intersection. The remaining strains also have a considerable number of specific genes although smaller than the core-genome shared between the six strains. Taken together, the *Burkholderia* specific genome is 48% larger than its counterpart in *Paraburkholderia* species.

### The global transcriptomic response to root exudates is strain-specific

In order to unravel bacterial genes involved in the preliminary steps of plant interaction, we stimulated the six *Burkholderia s.l.* strains with Nipponbare rice RE and compared their transcriptomic response to that of bacteria grown in VSG medium using three replicates by strain and condition (see Methods). A preliminary validation of the sequencing procedure confirmed that ribodepletion efficiency was > 70% for every strain and condition except for BvLMG10929 which only attained ∼50% efficiency (Supplementary Table S1). Nonetheless, high enough read numbers mapping against a database of CDSs were obtained for robust bioinformatic analyses (Supplementary Table S1). Furthermore, we verified that the exposition to RE significantly and consistently affected the bacteria’s transcriptomic responses throughout the replicates (Supplementary Figure S2).

At a quantitative level, the general response of bacteria to RE was strain-specific with large variations between strains. Besides BoABIP444, all strains display a majority of negatively regulated genes upon root exudate stimulation (Figure 4). For PspABIP659, the amount of down- and up-regulated genes is close to identical and represents the strongest response in terms of number of genes involved which is 8-fold stronger than for the weakest responder BrABIP441.

**Figure 4.**
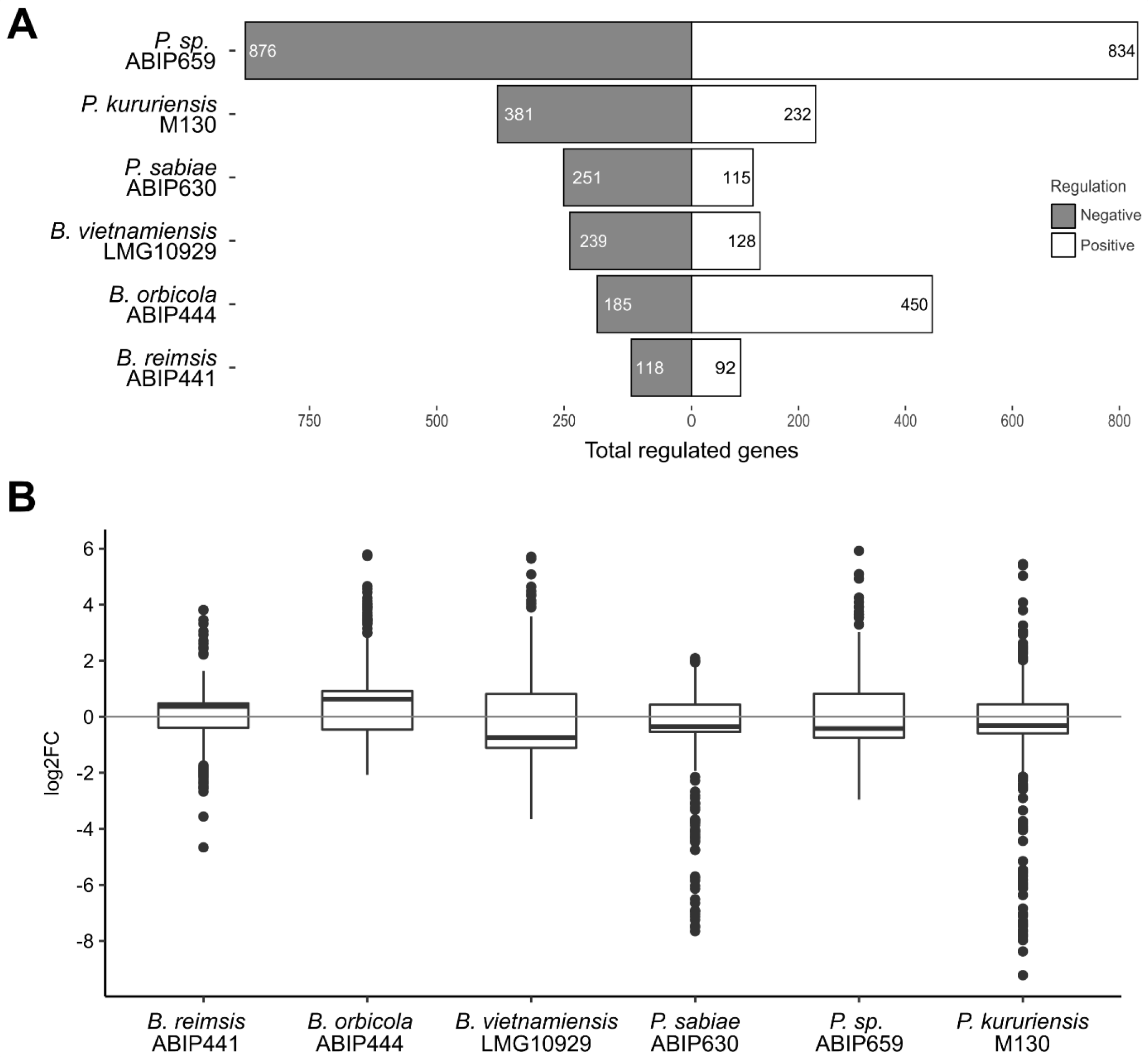
Transcriptomic response of *Burkholderia* and *Paraburkholderia* strains to RE. Genes that underwent a 1.5-fold and higher positive or negative regulation at a confidence level of adjusted pval ≤ 0.05 were considered to be significantly regulated upon stimulation with RE. The number of genes passing that threshold (A) and their regulation intensity on a log2 scale (B) are represented.

We further undertook an analysis of the conserved transcriptomic response across strains. Despite the level of genetic conservation between the six strains (Figure 3), the core-genome is not used in the primary steps of plant sensing as only two genes are DE in all six bacteria (Figure 5). One codes for an outer membrane porin (Table 2 & Supplementary Table S2) without further characterization. It is down-regulated in PsABIP630 and up-regulated in all remaining strains. The second encodes a TonB protein (Table 2 & Supplementary Table S2) which is associated with outer membrane receptors ExbB and ExbD. The differential expression of e*xbBD* is conserved in all strains besides BrABIP441 and follows the trend of the respective *tonB* regulation in each strain. It is up-regulated in PspABIP659 and BvLMG10929 and down-regulated in PsABIP630 and PkM130. If we exclude BrABIP441 from the analysis, a final gene is detected to be up-regulated in every strain, the Entner-Doudoroff pathway enzyme coding gene *edd* (Table 2 & Supplementary Table S2).

**Figure 5.**
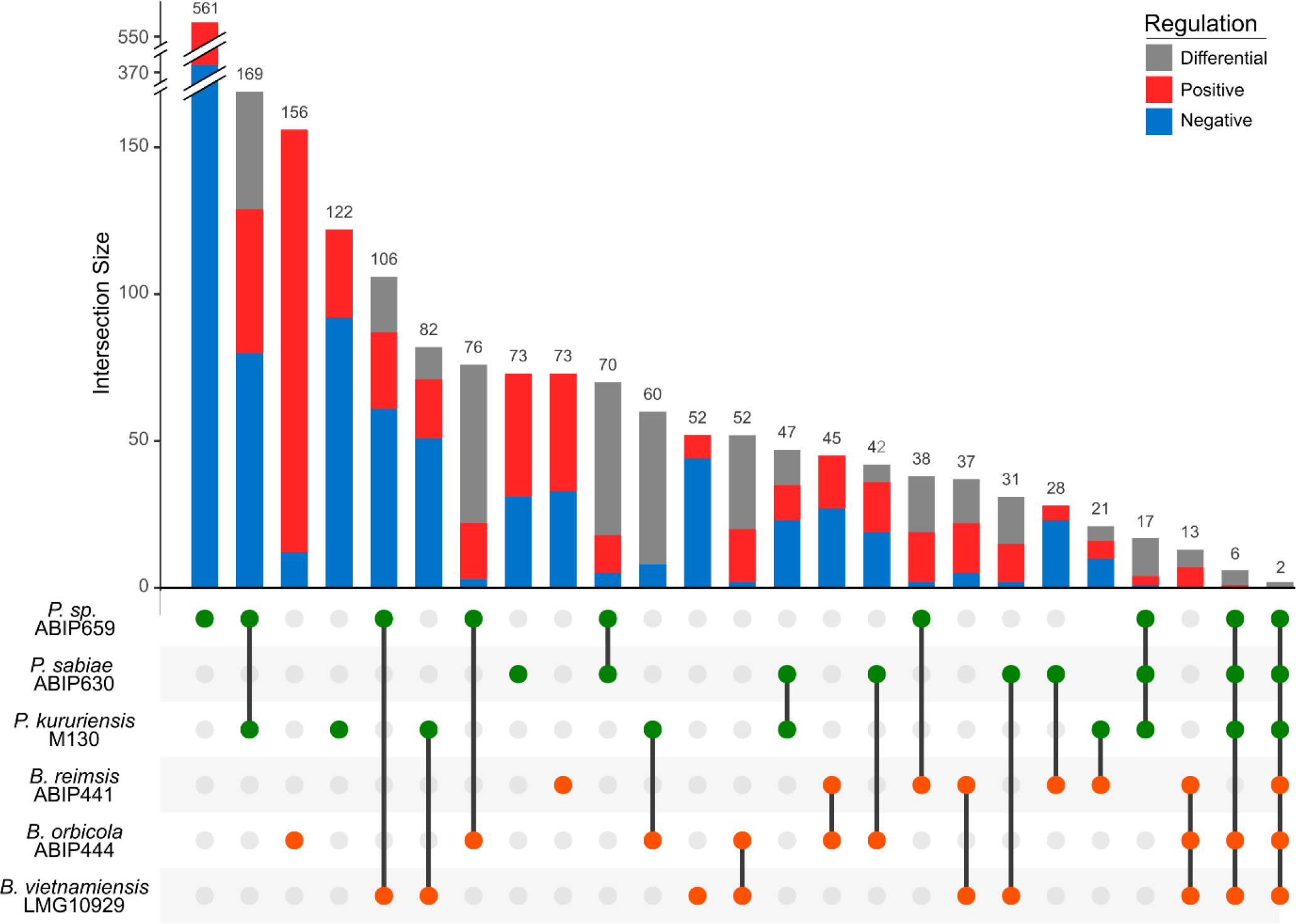
Specific and conserved regulation between strains upon RE stimulation. We calculated the conserved genome between different groups of strains to determine which genes are commonly regulated (either positively (red), negatively (blue) or differentially (grey) between each pair of strains, the *Burkholderia* and *Paraburkholderia* trios, all 6 strains and all 6 except BrABIP441. Single dots represent the regulation of strain-specific genes. For the PspABIP659 specific response, the histogram was reduced to fit with the figure (total regulated genes = 561, with a similar number of up and down genes)

**Table 2.**
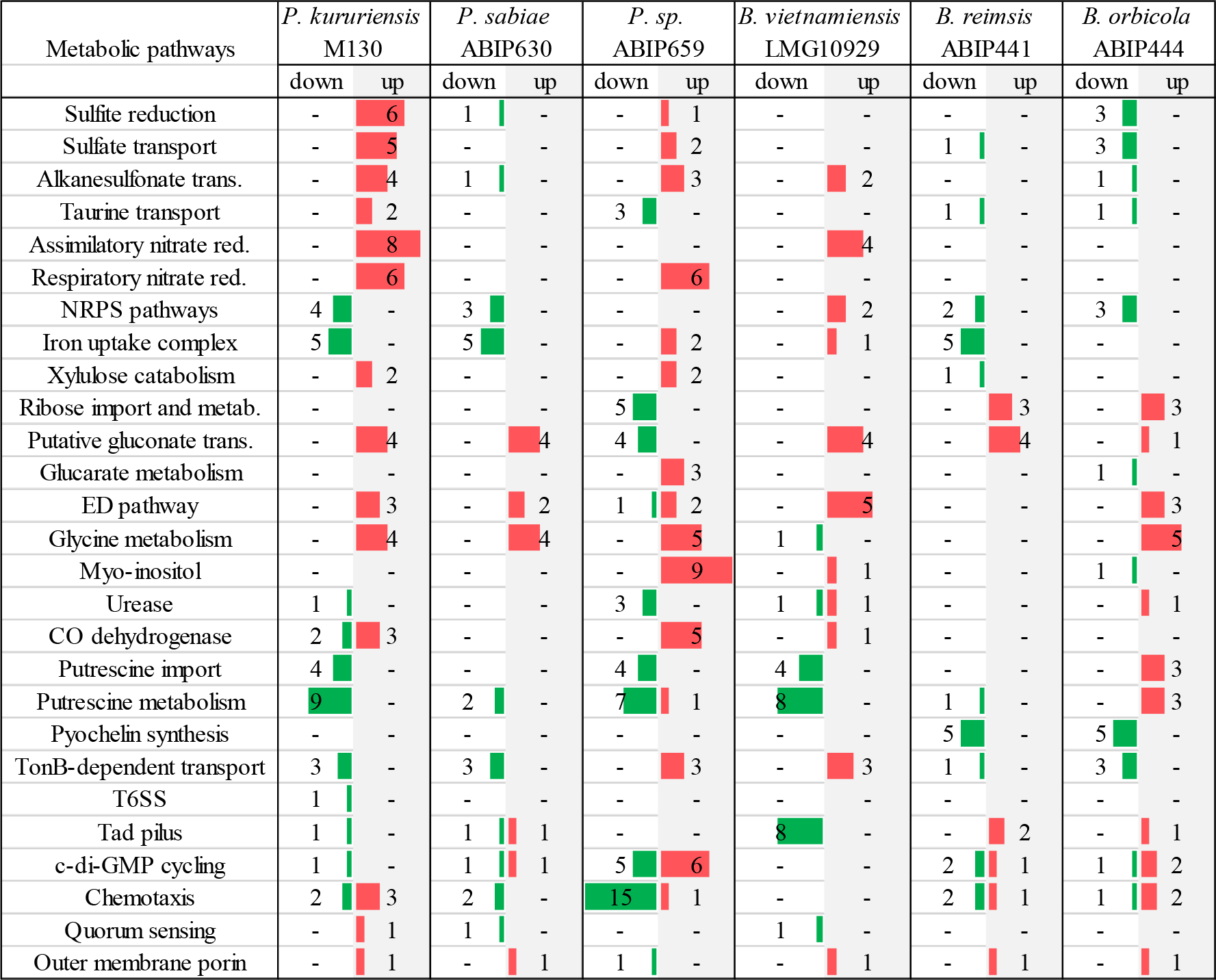
Differentially regulated gene categories in the six *Burkholderia s.l.* strains.

| Metabolic pathways | *P. kururiensis*<br>M130 |  | *P. sabiae*<br>ABIP630 |  | *P. sp.*<br>ABIP659 |  | *B. vietnamiensis*<br>LMG10929 |  | *B. reimsis*<br>ABIP441 |  | *B. orbicola*<br>ABIP444 |  |
| --- | --- | --- | --- | --- | --- | --- | --- | --- | --- | --- | --- | --- |
|  | down | up | down | up | down | up | down | up | down | up | down | up |
| Sulfite reduction | - | 6 | 1 | - | - | 1 | - | - | - | - | 3 | - |
| Sulfate transport | - | 5 | - | - | - | 2 | - | - | 1 | - | 3 | - |
| Alkanesulfonate trans. | - | 4 | 1 | - | - | 3 | - | 2 | - | - | 1 | - |
| Taurine transport | - | 2 | - | - | 3 | - | - | - | 1 | - | 1 | - |
| Assimilatory nitrate red. | - | 8 | - | - | - | - | - | 4 | - | - | - | - |
| Respiratory nitrate red. | - | 6 | - | - | - | 6 | - | - | - | - | - | - |
| NRPS pathways | 4 | - | 3 | - | - | - | - | 2 | 2 | - | 3 | - |
| Iron uptake complex | 5 | - | 5 | - | - | 2 | - | 1 | 5 | - | - | - |
| Xylulose catabolism | - | 2 | - | - | - | 2 | - | - | 1 | - | - | - |
| Ribose import and metab. | - | - | - | - | 5 | - | - | - | - | 3 | - | 3 |
| Putative gluconate trans. | - | 4 | - | 4 | 4 | - | - | 4 | - | 4 | - | 1 |
| Glucarate metabolism | - | - | - | - | - | 3 | - | - | - | - | 1 | - |
| ED pathway | - | 3 | - | 2 | 1 | 2 | - | 5 | - | - | - | 3 |
| Glycine metabolism | - | 4 | - | 4 | - | 5 | 1 | - | - | - | - | 5 |
| Myo-inositol | - | - | - | - | - | 9 | - | 1 | - | - | 1 | - |
| Urease | 1 | - | - | - | 3 | - | 1 | 1 | - | - | - | 1 |
| CO dehydrogenase | 2 | 3 | - | - | - | 5 | - | 1 | - | - | - | - |
| Putrescine import | 4 | - | - | - | 4 | - | 4 | - | - | - | - | 3 |
| Putrescine metabolism | 9 | - | 2 | - | 7 | 1 | 8 | - | 1 | - | - | 3 |
| Pyochelin synthesis | - | - | - | - | - | - | - | - | 5 | - | 5 | - |
| TonB-dependent transport | 3 | - | 3 | - | - | 3 | - | 3 | 1 | - | 3 | - |
| T6SS | 1 | - | - | - | - | - | - | - | - | - | - | - |
| Tad pilus | 1 | - | 1 | 1 | - | - | 8 | - | - | 2 | - | 1 |
| c-di-GMP cycling | 1 | - | 1 | 1 | 5 | 6 | - | - | 2 | 1 | 1 | 2 |
| Chemotaxis | 2 | 3 | 2 | - | 15 | 1 | - | - | 2 | 1 | 1 | 2 |
| Quorum sensing | - | 1 | 1 | - | - | - | 1 | - | - | - | - | - |
| Outer membrane porin | - | 1 | - | 1 | 1 | - | - | 1 | - | 1 | - | 1 |

Beside the intersection between PspABIP659 and PkM130, the levels of co-regulation are often higher in inter-generic than in intra-generic duos (Figure 5). The levels of conserved regulation within *Burkholderia* and *Paraburkholderia* trios is noticeably low and dominantly differential in *Paraburkholderia* (Figure 5).

### The transcriptomic response is mainly nutrient directed

The classification according to COG categories of the DE genes comforts the hypothesis that each strain has a dominantly individual response towards RE (Figure 6). Still, some trends can be inferred. In each strain, most DE genes involved in carbohydrate transport and metabolism are upregulated, indicating that the strains are taking advantage of the increased source of carbon. The reaction to other nutrients such as amino acids, lipids and ions is more disparate from strain to strain. Still, it appears that *Paraburkholderia* strains are globally more sensitive to the presence of lipids contained in RE.

**Figure 6.**
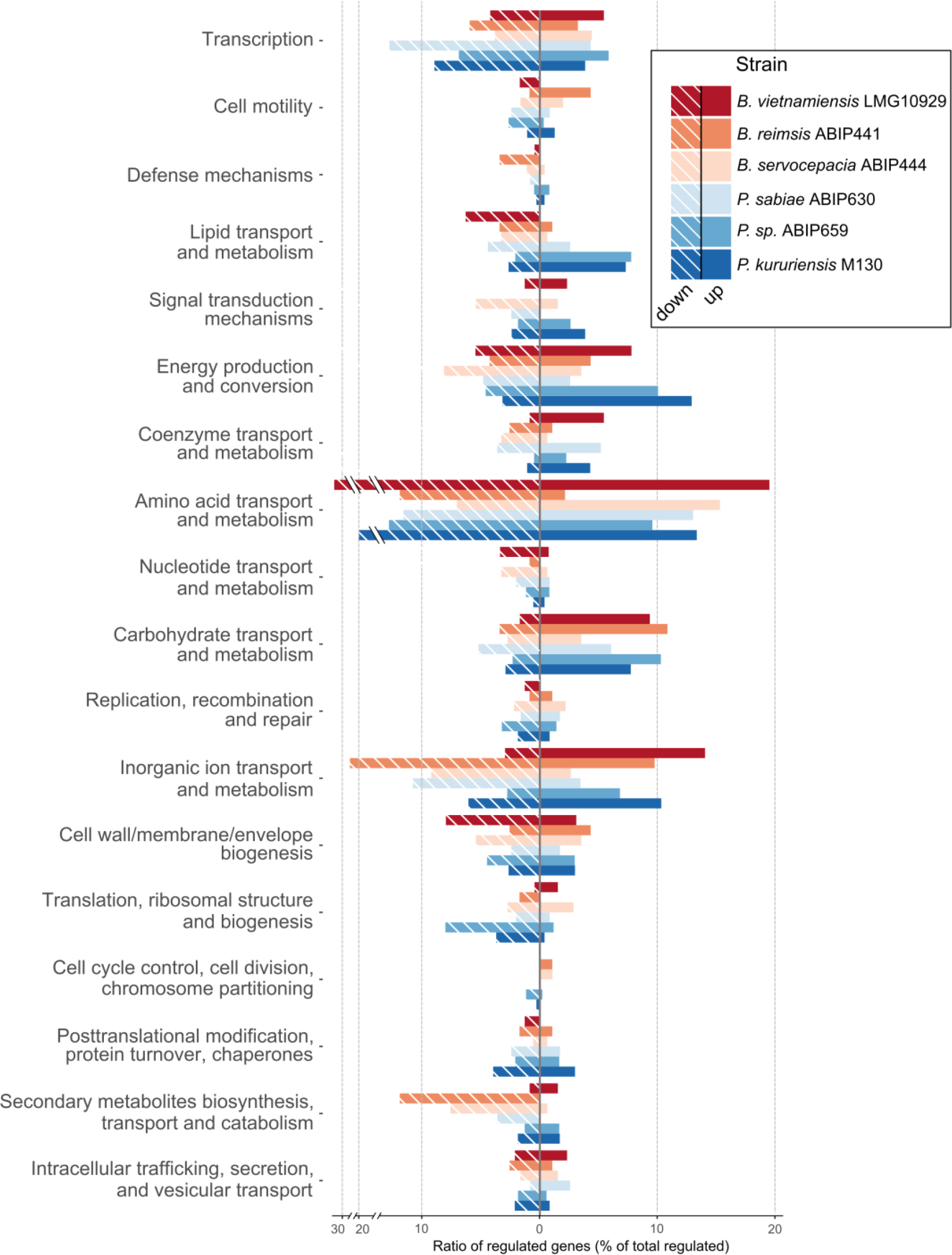
COG classification of the transcriptomic response of *Burkholderia* and ***Paraburkholderia* strains to RE.** The number of genes falling in each COG category is expressed as a percentage of the total amount of genes regulated in the corresponding strains (Figure 4A).

Our experimental approach used RE that were recovered from rice seedlings having grown for 5 days in a 6-fold diluted Hoagland nutritive medium (Supplementary Table S3). This experimental setting has been used before to assess the plant’s transcriptomic response elicited upon bacterial inoculation (King et al., 2019). However, diluted nutrients contained in this medium were assimilated to the recovered RE fraction. Thus, the bacterial response concerning elements contained in the hydroponic medium must be considered with care. As it was designed for plant nutrition the hydroponic medium does not contain any source of carbon. It does however hold macro-nutrients such as nitrogen, phosphorus and potassium. Additional elements eliciting a bacterial response are iron and sulfur. Still, the transcriptomic response to these nutrients varies depending on the bacterial strain (Table 2 & Supplementary Table S2).

Overall, we detected elements of the sulfite reductase complex (*cys*), the alkanesulfonate transport system (*ssuBCD*) and the taurine transporter *tauABC* which can be linked to sulfure nutrition and are DE only in *Paraburkholderia* strains. Elements linked to both assimilatory and respiratory nitrate transport and reduction were DE in several strains (Table 2 & Supplementary Table S2). Concerning iron nutrition, whole NRPS clusters, siderophore receptors and bacterioferritins are dominantly negatively regulated in most strains.

While nitrogen is a main constituent of the hydroponic medium, it is present as ammonium- conjugates and not nitrates. An increased nitrate metabolism was reported upon stimulation of PkM130 with rice macerate (Coutinho et al., 2015). The same study further reported the upregulation of a transporter complex involved in sulfate acquisition (ANSKv1_70577-81) which is also upregulated in the present study. Furthermore, the iron uptake complex is comparably downregulated in response to rice macerate (Coutinho et al., 2015) and what we observe with RE. These observations attest that the supposedly medium-induced regulations described above could in fact result from the composition of RE.

### Amino acid metabolism is activated upon exudate sensing

Several strains respond to the presence of amino acids from the exudates by downregulating the synthesis of dedicated transporters consistent with the strong adaptation of amino acid metabolism in response to RE which was repeatedly reported (Mark et al., 2005; Fan et al., 2012; Yi et al., 2017).. PkM130, BvLMG10929 and PspABIP659 all reduce the transcription of glutamate, aspartate and branched chain amino acids transporters. PspABIP659 further reacts in a similar fashion to the presence of arginine, lysine and histidine (Table 2 & Supplementary Table S2). The latter is also eliciting a response in PkM130. Upregulation of amino acid transporters was only observed for methionine in PkM130 and glycine in BoABIP444 (BCEN4_v1_340003).

Interestingly, an operon coding for a sarcosine oxidase (*soxBDAG*), a serine deaminase (*sdaB*) and an aldolase (*folB*) is up-regulated in all *Paraburkholderia* strains and in BoABIP444 but not in the remaining *Burkholderia* despite them bearing homologues of the concerned genes (Table 2 & Supplementary Table S2). Sarcosine is an intermediate in glycine synthesis which is generated by the sarcosine oxidase with hydrogen peroxide as a byproduct. The serine deaminase converts serine but also threonine into pyruvate and thus participates in gluconeogenesis. The FolB aldolase is an enzyme involved in one step of tetrahydrofolate synthesis, a cofactor used by the sarcosine oxidase.

### Exudates have a limited impact on chemotaxis and motility in our experimental design

Flagellar synthesis was not significantly impacted in our experiment. The chemotaxis pathway transducing elements *cheA*, *cheY* and *cheB* were equally devoid of regulation in all strains except *Psp*ABIP659 where *cheA*, *cheB* but also *cheW*, *cheD* and *motB* are negatively regulated (Table 2 & Supplementary Table S2). In other strains, chemotaxis elements present a more scattered regulation. The steadiness of genes involved in cell motility might be explained by their already high base level expression. The base mean of the flagellin component *fliC* reaches up to hundreds of thousands of reads depending on the strain. While, motility is usually activated upon root exudate sensing (Klonowska et al., 2018), in the present setting bacteria may have had already reached their induction maxima.

### Root exudate perception induces a shift in glycolysis strategies

The strains display numerous and specific reactions to the presence of carbohydrates from RE. Several strains positively regulate their dedicated metabolism to the presence of xylulose, ribose and glucarate (Table 2 & Supplementary Table S2).

A more conserved metabolic shift was however observed for glycolysis processes. In all strains except BrABIP441, genes involved in gluconate import and metabolism are up regulated. These enzymes are part of the Entner-Doudoroff (ED) pathway, an alternative to EMP glycolysis with similar costs and outcomes (Figure 7A). We previously demonstrated that ED pathway mutants of PkM130 and BvLMG10929 were significantly affected in their capacity to colonize rice roots (Wallner et al., 2022). The upregulation of the ED pathway by PkM130 and BvLMG10929 in the present study underlines its involvement in plant colonization as one of the early metabolic adaptations. Genes, downstream of the Entner-Doudoroff pathway, coding for a sugar ABC transporter are up-regulated in every strain except *Psp*ABIP659 where they are repressed.

**Figure 7.**
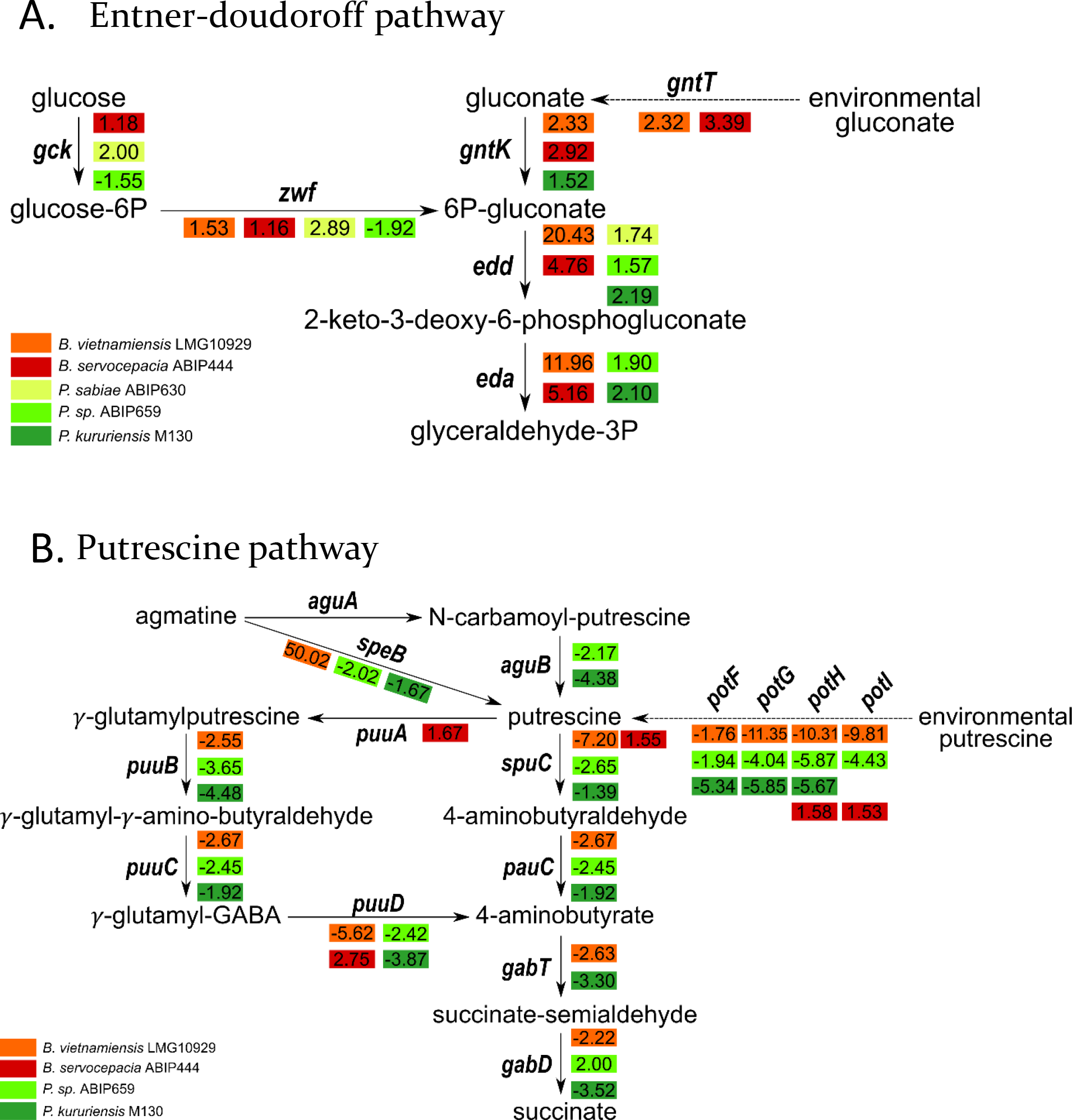
Regulation of Entner-Doudoroff and putrescine pathways upon root exudates stimulation. For each strain and pathway, the fold-change value is depicted next to the corresponding gene. (A) The Entner-Doudoroff pathway uses glucose as starting product and branches into the glycolysis pathway with glyceraldehyde-3P which is ultimately converted to pyruvate. No significant regulation was found in *B. reimsis* ABIP441. (B) Putrescine synthesis, catabolism and uptake pathways in *Burkholderia s.l.* strains. No significant regulation was found in *B. reimsis* ABIP441 and *P. sabiae* ABIP630.

### The perception of putrescine induces strain-specific adaptations

We paid particular attention to genetic elements that could indicate the onset of biofilm production as these structures are often involved in surface colonization. We noticed that RE were impacting the expression of genetic pathway involved in putrescine metabolism and cyclic diguanylate monophosphate (c-di-GMP) cycling. Upon stimulation with RE, we observe three different types of reactions to putrescine. PkM130, PspABIP659 and BvLMG10929 have a conserved response consisting in the downregulation of the previously mentioned genes. In PkM130 and PspABIP659, genes involved in putrescine formation from the precursor agmatine are also downregulated. Finally, BoABIP444 displays a fragmented regulation of the putrescine metabolism pathway but with systematic up-regulation of the respective genes (Table 2 & Figure 7B). The remaining strains, while they possess the according genes, do not adapt the pathways regulating the levels of intracellular putrescine.

### Cyclic di-GMP signaling is only activated in *Paraburkholderia* species

Intracellular c-di-GMP levels are regulated by diguanylate cyclases (DGC) for synthesis and diguanylate phosphodiesterases (PDE) for degradation. *Burkholderia* strains do not differentially express any of their genes involved in c-di-GMP signaling. This is coherent with what was previously observed through a Tn-seq approach in BvLMG10929 where no gene involved in c-di-GMP cycling was required for rice colonization (Wallner et al., 2022). For *Paraburkholderia* strains, PkM130 shows the most moderate response with the downregulation of a single DGC. PsABIP630 downregulates one DGC and upregulates one PDE. The strongest response is observed for PspABIP659 which differentially expresses eleven genes involved in c-di-GMP cycling. The majority of DGC are upregulated and the only two regulated PDE are repressed. Hence, there are different strategies at play for *Paraburkholderia* strains upon sensing of RE concerning the adaptation of their c-di-GMP signaling. PkM130 and PspABIP659 elevate the intra-cellular levels of the secondary messenger while PsABIP630 decreases them. Again, these results correlate with previous observations for PkM130 where a mutant involved in c-di-GMP cycling had a severe defect in rice colonization (Wallner et al., 2022).

## Discussion

In this study we analyzed the transcriptomic response to RE of six strains belonging to *Paraburkholderia* and *Burkholderia*. Given the ecologic background of both taxa we hypothesized that genera-dependent adaptation patterns could appear in the transcriptomic regulation upon sensing of rice RE. *Burkholderia* and *Paraburkholderia* strains share a close taxonomic proximity and thus a substantial level of orthologous genes. However, their strain-specific genomes are also of considerable size which is a premonitory sign of the disparate transcriptomic response displayed by the different strains upon chemical stimulation with RE. No categorizing trend could be inferred by the amount and intensity of regulated genes or by the function of regulated genes. Each strain displays a dominantly individual response and the genetic proximity between strains within a genus is not reflected in their transcriptomic response. Thus, in the present cases, the adaption to the habitat is independent from the genetic background.

Care was taken to expose the different bacteria to RE for a duration that was relative to their respective metabolic rate. The purpose was to obtain a synchronized response for reliable comparison. It is however conceivable that the six strains differently prioritize their responses which could result in an overall similar strategy that cannot be detected if sampled at a single time point. To avoid such biases induced by temporal shifts, a transcriptomic kinetic analysis would have been required. However, given the extent of differences observed here, we feel confident that there is indeed a strain-specific response to RE.

As the reported observations were made in a gnotobiotic setting, they are unlikely to reflect the full spectrum of bacterial adaptations in a complex natural environment. Still, species of *Burkholderia* s.l. were repeatedly described to be dominant in the rice rhizosphere which suggests that they can exist in communities where they exert the most influence beside their host (Ikeda et al., 2014; Yu et al., 2018).

Putrescine represents a long-overlooked metabolite in plant-bacteria interactions. While it has been identified as a virulence factor in *Ralstonia solanacearum* (Shi et al., 2019) it is also involved in the signalization between plant beneficial bacteria and their host (Liu et al., 2018). Putrescine is recognized as an important signaling molecule in plants where it is involved in various situations from organ development to biotic stress resilience (Tiburcio et al., 2014). Its presence in RE has been reported for tomato and rice (Kuiper et al., 2001; Suzuki et al., 2009) but has otherwise rarely been investigated. A study on the fitness determinants of *Pseudomonas sp*. WCS365 in the rhizosphere of wild type *Arabidopsis thaliana* and immunity impaired mutants demonstrated that the absence of putrescine catabolism genes (*spuC, pauC* and *gabT*) were highly increasing the bacteria’s fitness in the rhizosphere of WT but not immunocompromised plants (Liu et al., 2018). BvLMG10929 transposon mutants affected in putrescine metabolism and uptake were also detected to be less competitive during rice root colonization (Wallner et al., 2022). It is suggested that putrescine might act as a signaling molecule, informing the bacteria of the presence of a eukaryotic host. Responsive bacteria in turn promote biofilm formation which favors attachment to the root. This regulatory system must be extremely fine-tuned as to not trigger plant defenses by an excess production of biofilms and other physiological changes subsequent to putrescine signalization. In the transposon sequencing assay on *Pseudomonas* sp. WCS365, genes involved in putrescine uptake and synthesis were found to be negative fitness regulators (Liu et al., 2018). Consistently, PkM130, PspABIP659 and BvLMG10929 downregulate the according gene homologues upon exudate sensing. Thus, we could hypothesize that these strains take advantage of putrescine to regulate their metabolism and physiology towards a controlled surface colonization of roots.

C-di-GMP is another signal molecule positively regulating biofilm formation when accumulated (Romling et al., 2013). The regulation of enzymes involved in its metabolism can therefore be key for a successful colonization of plant roots and evasion of host defenses. We previously demonstrated the involvement of a c-di-GMP cycling enzyme in the root colonization process of rice by PkM130 (Wallner et al., 2022). Inversely, in BvLMG10929 several genes involved in c-di-GMP cycling were identified as being obstructive to a successful root colonization (Wallner et al., 2022). Recently, the importance of c-di-GMP in biofilm production and subsequent virulence on rice was demonstrated for *Burkholderia glumae* BGR1 (Kwak et al., 2020). The human opportunist *B. cenocepacia*, which is closely related to *B. orbicola,* received much attention for the role played by c-di-GMP signalization in virulence by regulation of biofilm formation and motility (Richter et al., 2019). Similarly, the absence of the PDE-receptor coding gene *rpfR* in *Burkholderia lata* SK875, reduces the strain’s virulence and increase its biofilm production (Jung et al., 2017). RpfR is a receptor for the *Burkholderia* specific quorum sensing signal BDSF which implies a connection between c-di-GMP and cell-to-cell signaling (Schmid et al., 2017). Further studies linked the production of biofilms with the sensing of nitrate and temperature which are relying on a subsequent c-di-GMP dependent signaling pathway in *B. pseudomallei* (Mangalea et al., 2017; Plumley et al., 2017). The maize endophyte *Burkholderia gladioli* 3A12 displays c-di-GMP dependent signaling for biocontrol purposes against fungal pathogens (Shehata and Raizada, 2017). Although root exudate stimulation had no direct impact on the regulation of flagellum-related genes, the regulation of genes involved in putrescine uptake and metabolism as well as those involved in c-di-GMP signaling indicate an indirect and intricate regulation of motility (Liu et al., 2018; Richter et al., 2019). This is coherent with the supposition that bacteria evade host defenses by reducing the production of the common flagellin MAMPs. This behavior was described upon exudate stimulation in *Pseudomonas aeruginosa* PA01, *Azoarcus sp.* BH72 and *P. kururiensis* M130 (Mark et al., 2005; Shidore et al., 2012; Coutinho et al., 2015).

Until recently, the ED pathway had never been reported for its activation at any stage of plant colonization. The difference with the more classic EMP-glycolysis pathway is the generation of NAD(P)H cofactor during ED-glycolysis. This cofactor can be essential for enzymes that play a decisive role in rhizospheric adaptation as it is the case for thioredoxins, involved in oxidative stress tolerance (Cumming et al., 2004; Chavarría et al., 2013). The *edd* gene, coding for the NAD(P)H-generating enzyme, is amongst the only genes that are differentially regulated in all six strains. In a separate study, we identified *edd* as being critical for a successful interaction of BvLMG10929 and PkM130 with rice roots as its absence led to a significant decrease in the bacteria’s colonization capacities (Wallner et al., 2022).

Using six different strains from the *Burkholderia* and *Paraburkholderia* genera, we could find no discerning pattern in their early response to the perception of plants through RE, suggesting that there is a strong intra-generic diversity. Thus, instead of analyzing one *Burkholderia* and one *Paraburkholderia* group composed of three species each, we rather have six distinct strains that independently underline the importance of certain pathways in plant-bacteria interactions. While some, such as c-di-GMP signaling were already recognized, the implication of putrescine and the ED-pathway should receive an increased attention and especially in later steps of plant colonization.

## Methods

### Plant cultivation and exudate collection

*Oryza sativa* ssp. *japonica* cv. Nipponbare seeds were sterilized, germinated and cultivated in a sterile, hydroponic system as described before (King et al., 2019). Five seedlings were planted per system and transferred to a growth chamber (28°C; 16 h light; 8 h dark). After 3 days of growth, the hydroponic medium was collected using a sterile graduated pipette and filtered using sterile Millipore membranes to remove residual perlite sand. The resulting medium was frozen at -80°C and lyophilized. Dry RE were resuspended in sterile distilled water at an 80x concentration and stored at -20°C until use.

For plant growth measurements, rice seedlings were sterilized and germinated as before and transferred to pots containing 400 mL of a 7/2/1 mix of sterile sand (0.6-1.6 mm)/ perlite/vermiculite. Five pots containing four rice seedlings were used per condition.

### Bacterial strains and cultivation

*Burkholderia reimsis* ABIP441, *Paraburkholderia* sp. ABIP659 and *Paraburkholderia sabiae* ABIP630 were isolated from healthy *Oryza sativa* subsp. *japonica* roots. ABIP441 and ABIP630 were isolated from rice roots collected from a field in Cameroon and ABIP659 from a field in Vietnam. Additionally, we used the well characterized rice growth promoting species *Burkholderia vietnamiensis* LMG10929 and *Paraburkholderia kururiensis* M130 as well as the recently proposed *Burkholderia orbicola* sp. nov. (previously named *B. servocepacia*) ABIP444 (Wallner et al., 2019; Morales-Ruíz et al., 2022).

All bacterial strains were cultured as follows: Glycerol stocks (20%) of bacterial cells conserved at -80°C were plated on Luria’s low salt LB (LBm; Sigma-Aldrich) agar plates and incubated for 72 h at 28°C. Single colonies were used to inoculate 10 mL of LBm broth in 50 mL Falcon tubes and incubated for various amounts of time allowing the different strains to reach an OD_600_=1.0. For DNA extraction purposes, cells were used at this stage.

### Plant inoculation assays

For colonization assays, bacterial strains were grown overnight in liquid LBm as described above. Bacterial density was then adjusted to 1.10^8^ cfu.mL^-1^ with regard to the relative amount of cells at OD_600_=1.0 for each strain. In a sterile setting, 5-day old rice seedlings, prepared as described above, were inoculated with 1 mL of the bacterial solution at the base of each plant. Two systems (10 plants total) were prepared for each condition and grown for 14 days (28°C; 16 h light; 8 h dark). Five plants were randomly harvested per condition as biological replicates. The entire root system was sampled and transferred to a screw cap tube containing 1 mL of sterile water and a sterile ceramic bead. Roots were weighted and then pulverized using a Fastprep-24 (MPbio) at 6 m.s^-1^ for 40 seconds. A serial dilution of the resulting solution was spotted out on square LBm plates and incubated at 28°C for 48 hours.

For plant growth measurements overnight broth bacterial cultures in LBm were pelleted by centrifugation, resuspended in distilled water and adjusted to OD_600_=1.0. CFU measurements at this OD are reported in the legend of supplementary Figure S1. Twenty plants grown in a sand/perlite/vermiculite mix were inoculated with 1 ml of bacterial suspension per plant, 5 days after seedling transfer to pots. Plants were grown in a growth chamber (80% humidity, 28°C; 16 h light; 8 h dark) for 50 days before sampling, and fertilized with a 50% Hoagland solution, twice a week. The length of the longest leaf as well as the root’s and leave’s dry weights were measured for each plant. Statistics were performed in R using packages rstatix, multcompView and ggplot2 (Wickham, 2016; Kassambara, 2020; Graves et al., 2022) and differences were estimated with a pairwise Wilcoxon non-parametric test. R Script used to do statistics and produce Supplementary Figure S1 is accessible on Zenodo (https://doi.org/10.5281/zenodo.7447236).

### Exudate stimulation of bacterial cells

For RNA extraction, cells were prepared as detailed above but using Vincent Succinate Glutamate (VSG) growth medium (Vincent, 1970). The pre-culture was then used to inoculate 50 mL VSG in 250 mL Erlenmeyer flasks at a final OD_600_=0.075. After 1 h of agitation at 150 rpm and 28°C, 800 µL of concentrated root exudate solution (62-fold) was added to obtain a final exudate concentration that was equivalent to that found in the hydroponic growth system. Cells were then grown until they reached mid-exponential phase. The volume required for 10^9^ cells was collected and immediately transferred to a 15mL Falcon tube containing 95% ethanol and 5% phenol. Collected cells were centrifuged at 4000 rpm for 15 minutes. The supernatant was discarded and the pellet was used to extract total RNA using the RiboPure RNA purification kit for bacteria (ThermoFisher). After extraction, the sample were subjected to a TURBO DNase (ThermoFisher) treatment to remove any residual DNA. Extraction efficiency was qualified using the 2100 Bioanalyzer system with an RNA Nano Chip (Agilent Technologies).

### Ribodepletion and RNA sequencing

Ribosomic RNA (rRNA) was removed from total RNA extracts following the guidelines of Roux et al., 2018. Briefly, we generated probes targeting 5S, 16S and 23S rRNA incorporating a biotin complex in 5’ and locked nucleic acids (LNAs) for improved specificity towards rRNA of the burkholderia group (Supplementary Table S4). An *in silico* approach validated that these probes are universal to all *Burkholderia s.l.* species and conserve specificity in the neighboring taxa *Ralstonia* and *Cupriavidus.* 2 µL of probes (50 µM) were used in combination with reagents from the RiboMinus Transcriptome Isolation Kit, bacteria (ThermoFisher). We used 2 µg of total RNA and 71 µL of magnetic beads. Prior to use, magnetic beads were washed twice in nuclease free water and once in 2x SSC buffer and resuspended in a final volume of 31 µL hybridization buffer. Ribodepletion efficiency was validated using the 2100 Bioanalyzer system (Agilent Technologies) with an RNA Nano Chip (Supplementary Table S1).

RNA sequencing was performed at the high throughout sequencing platform at I2BC (CNRS, Gif-sur-Yvette, France) using Nextseq500 (Illumina) technology with a Nextseq 500/550 High Output Kit v2 (Illumina) and 75 sequencing cycles (single read, 75 bp) on ribodepleted RNA samples originating from three biological replicates per strain and per growth condition. De-multiplexing, adapter trimming and quality controls were performed by the platform using bcl2fastq2-2.18.12, Cutadapt 1.15 and FastQC v0.11.5 respectively (Andrew, 2010; Martin, 2011). Raw reads and biosamples are accessible under Bioproject PRJEB42574 / ERP126457 at the European Nucleotide Archive (https://www.ebi.ac.uk/ena/browser/view/PRJEB42574).

Reads were mapped to the reference genomes using the CLC workbench and differential gene expression was inferred using the DESeq2 package (Love et al., 2014). Low-count genes were removed at a threshold of 5 reads and an independent hypothesis weighting was applied. One replicate had to be removed from the analysis for *B. reimsis* ABIP441 as the sequencing depth was insufficient to allow robust analyses (Supplementary Figure S2). Genes that had a positive or negative differential expression superior to 1.5-fold at pval < 0.05 were taken into consideration.

### DNA extraction and genome sequencing

The genomic DNA of ABIP441, ABIP630 and ABIP659 was extracted using the standard operating procedure of the JGI as described before (Wallner et al., 2019). A genomic library with average insert size of 350 bp was prepared for sequencing using a TruSeq Nano DNA library preparation kit (Illumina, Inc., San Diego, CA, USA). Paired-end 2 x 125 nt sequencing was performed by the MGX platform (CNRS, Montpellier, France) using a HiSeq 2500 (Illumina), generating 19,127,584 raw read pairs for ABIP441, 12,198,848 for ABIP630 and 16,208,618 for ABIP659. Reads with overlapping sequence were assembled using CLC Genomics Workbench version 7.04 resulting in 311 contigs ranging from 506 bp to 27.4 kbp for ABIP441, 84 contigs ranging from 811 bp to 424.5 kbp for ABIP630 and 122 contigs ranging from 1162 bp to 173.7 kbp for ABIP659.

The genome quality was assessed using BUSCO v3 for the detection of mandatory single-copy orthologues (Simão et al., 2015). Genome completeness was estimated at 97.0%, 99.0% and 97.8% for ABIP441, ABIP630 and ABIP659 respectively. Genbank accession numbers for the three new genomes are GCA_914492715 (ABIP441), GCA_914484815 (ABIP630), GCA_914484835 (ABIP659), under Bioproject PRJEB42536 (https://www.ncbi.nlm.nih.gov/bioproject/).

### Comparative genomics procedures

The genomic similarity was assessed using the Genome clustering tool of the Microscope platform https://mage.genoscope.cns.fr/microscope/ (Vallenet et al., 2020). It is based on Mash, a software that computes distances between two genomes (Ondov et al., 2016). From all the pairwise distances of the genomes set, a tree is constructed dynamically using the neighbor-joining javascript package (Simonsen et al., 2008). The clustering annotations are computed from all-pairs distances ≤ 0.06 (or 94% ANI), corresponding to the ANI standard to define a species group. The clustering has been computed using the Louvain Community Detection. Average Nucleotide identities were inferred using the ANI calculator tool (https://www.ezbiocloud.net/tools/ani).

Core genome compositions were calculated using the Phyloprofile exploration tool implemented in the MicroScope microbial genome annotation and analysis platform (Vallenet et al., 2020). Homology constraints were set at minLrap ≥ 0.8 and identity ≥ 50%. To establish a list of specific genes for comparative genomics purposes, exclusive comparisons were performed for the 63 possible combinations between the 6 strains where the genomes not used in the comparison were systematically use as negative filter at the same homology constraints as above. To establish a list of co-regulated genes between the six species, non-exclusive comparisons were performed for each pairwise combination as well as for the *Burkholderia* and *Paraburkholderia* trios and the combination of all six species.

