## Supplementary Table S1 for "Comparative genomics and transcriptomic response to root exudates of six rice root-associated *Burkholderia sensu lato* species"

**Supplementary Table S1**: High throughput sequencing results of ribodepleted RNA.

|  | *B. reimsis* ABIP441 | *B. orbicola* ABIP444 | *B. vietnamiensis* LMG109129 | *P. sabiae* ABIP630 | *P. sp.* ABIP659 | *P. kururiensis* M130 |
| --- | --- | --- | --- | --- | --- | --- |
| **Control** |  |  |  |  |  |  |
| Mean reads | 31,344,469 | 35,168,205 | 31,057,775 | 33,202,131 | 29,595,520 | 30,761,298 |
| Mapped unique reads | 24,652,795 | 26,458,901 | 16,091,391 | 26,845,561 | 26,609,142 | 22,261,009 |
| Ribodep. efficiency | 79% | 75% | 52% | 81% | 90% | 72% |
| **Exudates** |  |  |  |  |  |  |
| Mean reads | 34,952,561 | 36,054,658 | 31,611,613 | 32,836,389 | 29,992,068 | 38,333,436 |
| Mapped unique reads | 26,797,404 | 28,614,667 | 16,983,325 | 26,276,026 | 25,420,310 | 28,390,557 |
| Ribodepletion efficiency | 77% | 79% | 54% | 80% | 85% | 74% |
| Ribodepletion efficiency was inferred through direct mapping of reads against a database of ribosomal RNA.  Mapped unique reads were inferred through direct mapping against a database of CDSs. | | | | | | |
