## Supplementary Table S2 for "Comparative genomics and transcriptomic response to root exudates of six rice root-associated *Burkholderia sensu lato* species"

**Supplementary Table S2:** Culture media compositions.

| **Component** | **VSG** | **Hoagland** |
| --- | --- | --- |
| (NH_4_)_2_SO_4_ |  | 0.5 mM |
| (NH_4_)_6_Mo_7_O_24_ |  | 0.16 µM |
| Nicotinic acid | 0.406 µM |  |
| Biotine | 2.0 µM |  |
| Ca(NO_3_)_2_ |  | 1.2 mM |
| CaCl_2_ | 0.45 mM |  |
| CaSiO_3_ |  | 1.7 mM |
| CuSO_4_ |  | 0.8 µM |
| EDTA |  | 90.0 µM |
| FeCl_3_ | 37 µM |  |
| FeSO_4_ |  | 90.0 µM |
| H_3_BO_3_ |  | 22.6 µM |
| K2HPO_4_ | 11.2 mM |  |
| KH_2_PO_4_ | 14.4 mM | 0.4 mM |
| KNO_3_ |  | 0.7 mM |
| MgSO_4_ | 1.0 mM | 1.6 mM |
| MnSO_4_ |  | 10.0 µM |
| *Myo-inositol* | 2,78 µM |  |
| Panthothènate Ca | 0.193 µM |  |
| Pyridoxine | 0.296 µM |  |
| Sodium glutamate | 5.8 mM |  |
| Sodium succinate | 12.1 mM |  |
| Thiamine | 0.188 µM |  |
| ZnSO_4_ |  | 0.7 µM |
