## Supplementary Table S4 for "Comparative genomics and transcriptomic response to root exudates of six rice root-associated *Burkholderia sensu lato* species"

**Supplementary Table S4**: List of custom probes used for rRNA depletion.

| Target | # | Sequence (5'-3') |
| --- | --- | --- |
| 5S | 1 | GAG T+CG TT+T CAC GGT +CCT GT+T CG |
|  | 2 | GCC T+GA CGA TTA +CCT A+CT T+TC AC |
| 16S | 1 | ATG CGG TAT +TAA T+CC GG+C TT+T CG |
|  | 2 | TTC T+TC A+CA CA+C GCG G+CA TTG CT |
|  | 3 | CCG G+GG ATT TCA +CAT +CGG T+CT TA |
|  | 4 | GTT T+TA AT+C TTG CGA +CCG +TAC TC |
|  | 5 | TGT GA+C GGG +CGG TGT G+TA CA+A GA |
| 23S | 1 | TTT T+CG CA+G GCT A+CC GCG T+CC TT |
|  | 2 | CCC TCA +CGG TA+C TGG T+TC A+CT AT |
|  | 3 | CCA TG+G ATA +GAT CA+C CTG GT+T TC |
|  | 4 | CTC TTG A+CA C+CG GA+C GT+T AGC AC |
|  | 5 | TTT CA+C CC+C CT+T TAT CG+C TAC TC |
|  | 6 | TCA CAC C+CT ATA +CGT C+CA CT+T TC |
|  | 7 | CCA GAG +TAC +CTT T+TA T+CC GTT GA |
|  | 8 | ATC TCT T+CC GAA +CAT AG+C TA+C CC |
| Different quantities of probes were generated proportionally to the size of the targeted ribosomal sequence. LNAs are preceded by a (+). | | |
