## Supplementary Figure S1 for "Comparative genomics and transcriptomic response to root exudates of six rice root-associated *Burkholderia sensu lato* species"

**Supplementary Figure S1** : Impact of bacterial inoculation on rice at 50 days post -inoculation. Twenty plants were inoculated per condition with 1 ml of an OD_600_= 0.1 culture, corresponding to 8.9 x 10^7^ for BvLMG10929, 6.2 x 10^7^for Br441, 9.9 x 10^7^ for Bo444, 1.5 x 10^6^ for Ps630, 7 x 10^7^ for Psp659, 1 x 10^8^ for PkM130. Letters on top of box plots indicate results of non-paramteric test (Kruskall-Wallis at p≤0.05).

.

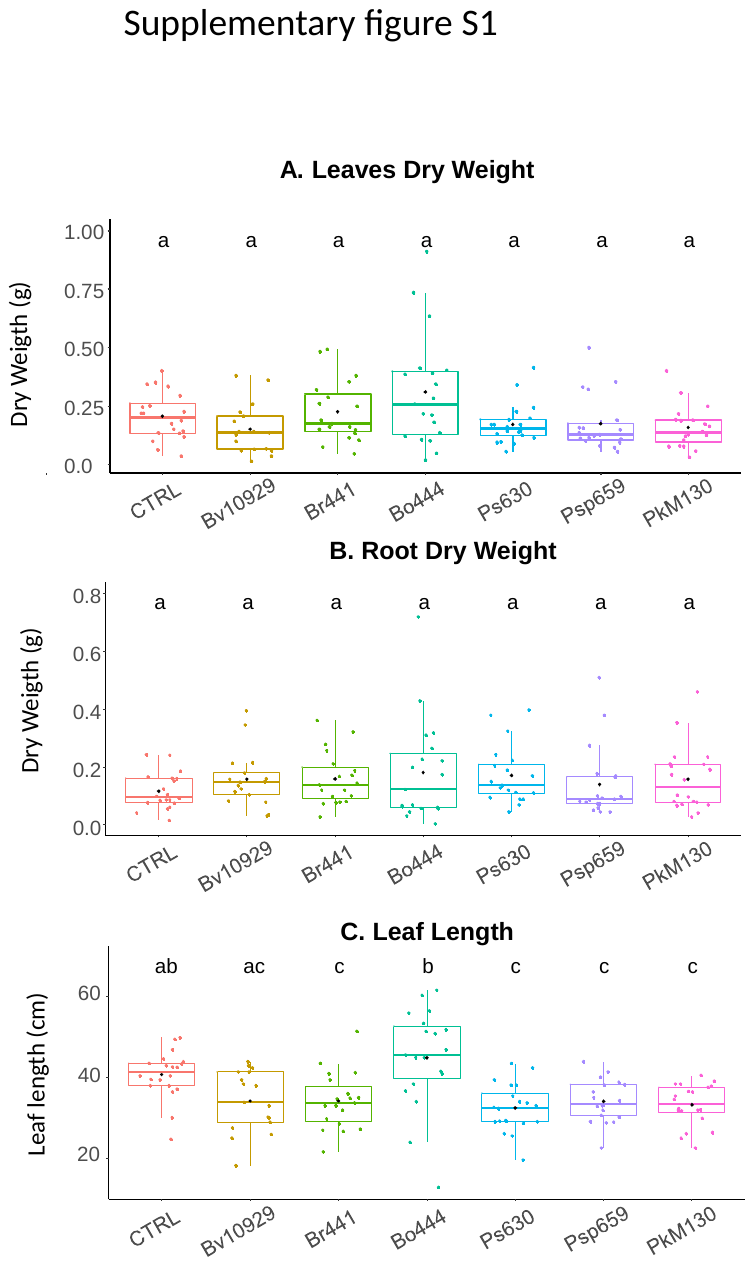
