## Supplementary Figure S2 for "Comparative genomics and transcriptomic response to root exudates of six rice root-associated *Burkholderia sensu lato* species"

**Supplementary Figure S2** : Variance analysis between transcriptome replicates. For ABIP441, one replicate of the exudate group failed to provide enough reads and was thus removed from the analysis

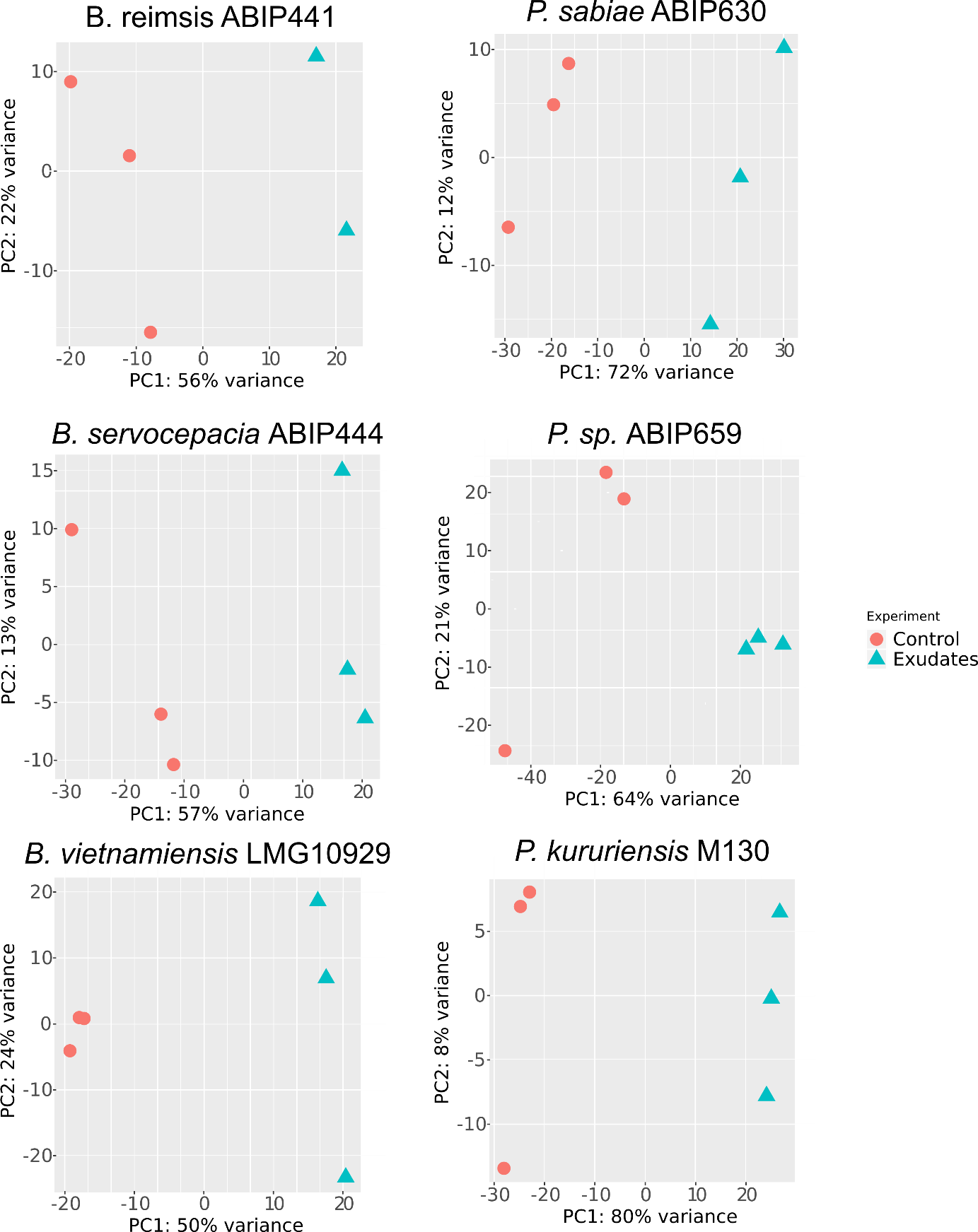
